# *Achatina fulica* Haemocyanin-Derived Peptides as Novel Antimicrobial Agents

**DOI:** 10.1101/2023.08.15.553437

**Authors:** Andrés Pereira, Libardo Suarez, Tanya Roman, Fanny Guzmán, Bladimiro Rincón-Orozco, Leidy Sierra, William Hidalgo

## Abstract

Anteriorly we found Haemocyanin-derived peptides in semi-purified fractions of mucus secretion of *Achatina fulica* that showed an inhibitory effect on *S. aureus* strains. Here, we applied *in silico* rational design strategy to obtain new potential Antimicrobial Peptides (AMPs) from *A. fulica* haemocyanin-derived peptides (AfH). Designed peptides were chemically synthetized using the Fmoc strategy and antimicrobial activity against *E. coli* and *S. aureus* strains was investigated using the broth microdilution method. Cytotoxic activity on Vero, HaCat, and human erythrocytes cells was also evaluated. The results showed that 15-residue synthetic peptides, alpha-helical and cationic, were those with the highest biological activity against Gram positive strains, with Minimum Inhibitory Concentrations (MIC) in the range of 20 to 30 µM. The positive selectivity index suggests a higher selectivity mainly on the microorganisms evaluated and not on eukaryotic cells. In this study, *A. fulica* hemocyanin turned out to be an appropriate protein model for the rational design of AMPs against bacteria of public health importance. Additional studies are required to evaluate the activity of the peptides on Gram negative bacteria.

## Introduction

Searching new compounds with antimicrobial activity is crucial in combating antimicrobial resistance. To discover innovative solutions, scientists have shifted their focus to invertebrates, particularly Phyla Mollusca. These organisms offer abundant sources of pharmacologically active compounds present in their haemolymph and mucus secretions (Sperstad et al., 2011; Tassanakajon et al., 2015). Exploring these natural reservoirs could provide valuable insights and potential antimicrobial agents in the ongoing fight against resistant pathogens (Neill, 2014; WHO, 2014).

Gastropods, including snails and slugs, represent the most diverse class of molluscs. Among terrestrial gastropods, only a handful of species have been studied for their antimicrobial properties, primarily focusing on mucous secretions. These species include *Achatina fulica* (Alogna, 2017; Cho et al., 2019; Daud et al., 2018; Vieira et al., 2004; Zhong et al., 2013), *Cornu aspersum* (Dolashki et al., 2018, 2020; Kostadinova et al., 2018; Velkova et al., 2018), *Archachatina marginata* (Adikwu, 2005; Kenechukwu et al., 2013), *Helix pomatia* (Greistorfer et al., 2017), *Hemiplecta distinta* (Noothuan et al., 2021), and *Arion ater* (Cottrell et al., 1994). The choice of the species for investigation depends on their availability and economic significance.

*A. fulica* is a gastropod native to eastern Africa, and it has been reported in more than 60 countries across five continents. Its invasive nature has led to its inclusion in the list of the 100 World’s Worst Invasive Alien Species, primarily due to its detrimental impact on agriculture, ecosystems, human health, and the economy (Lowe et al., 2004). Despite numerous studies conducted on the mucus of this snail, which have revealed various biological activities such as the inhibition of microbial growth, inflammatory processes, and wound healing properties, there remains limited information available regarding antimicrobial compounds derived from *A. fulica* and terrestrial gastropods in general (Ghosh et al., 2010; Iguchi et al., 1982; Zhong et al., 2013).

The antibacterial effect of *A. fulica* mucus has been attributed to the presence of proteins and AMPs. However, apart from the well-characterized L-amino acid oxidase Achacin (59 kDa) and the cysteine-rich AMP mytimacin-AF (9.7 kDa), only a few antimicrobial compounds from *A. fulica* have been fully identified and characterized (Obara et al., 1992; Zhong et al., 2013). This scarcity of comprehensive understanding underscores the need for further exploration and investigation into the antimicrobial properties of *A. fulica* and its potential as a source of novel therapeutic agents against bacterial infections. Expanding our knowledge in this area holds great promise for addressing the urgent challenge posed by antibiotic-resistant bacteria and discovering alternative antimicrobial strategies.

Most of the mature AMPs arise from the intracellular processing of larger precursor proteins by post-translational modifications. Some studies suggest that biologically active proteins such as hemocyanin (Lee et al., 2003a) and haemoglobin (Ullal et al., 2008), may be sources of AMPs.

Hemocyanin (Hc) is an oxygen-carrying protein that only has been reported in the hemolymph of some cephalopods (molluscs), crustaceans, and arachnids (arthropods). Hc-derived peptides with antimicrobial properties have been reported in shrimp (Monteiro et al., 2020; Petit et al., 2016; Zhan et al., 2019), crayfish (Lee et al., 2003b), and spiders (Riciluca et al., 2012). The bibliographic review indicates that the first AMPs derived from mollusc hemocyanin were recorded until 2015 (Zhuang et al., 2015).

This study focuses on investigating *A. fulica* hemocyanin as a model protein for the design of novel peptides with antimicrobial potential. The hypothesis underlying is that rational *in silico* design of peptides derived from *A. fulica* hemocyanin can yield novel AMPs with the potential to combat bacteria of significant importance in public health, including Methicillin-Resistant *Staphylococcus aureus* (MRSA) and *Escherichia coli* O157:H7.

To achieve this, we conducted the design, synthesis, physicochemical characterization, and evaluation of synthetic peptides derived from *A. fulica* hemocyanin against *Staphylococcus aureus* and *Escherichia coli* strains that are resistant to conventional antibiotics such as methicillin and oxacillin. Furthermore, we assessed the cytotoxic and haemolytic activities of the peptides exhibiting the most promising biological activity.

This study contributes to the growing body of knowledge on the development of alternative antimicrobial strategies to address the challenges posed by antibiotic-resistant bacteria, offering potential solutions in the fight against multidrug-resistant pathogens.

## Materials and Methods

### Mucus collection

The specimens of *A. fulica* identified by morphological characteristics were collected in Floridablanca, Santander, Colombia. Mucus secretion was obtained by direct stimulation on the foot with 9V electric current at intervals of 30 to 60 seconds, according to the protocol stablished in (Pereira et al., 2016).

### Peptide design

Sequences of hemocyanin-derived peptides obtained from the mucus secretion of *A. fulica* (Suárez et al., 2021) were used as sources to design new analogous peptides with high antimicrobial potential. To archive this, a *in silico* rational design strategy was conducted using bioinformatics tools as follows:

Peptide sequences were aligned with the BLASTp tool to obtain the regions of the hemocyanin of terrestrial gastropods with identity greater than 80% with the peptides obtained from the mucous secretion of *A. fulica*. Clustal Omega multiple sequence alignment method was used to find the conserved residues in peptide sequences. Alignment with patterns of different families of AMPs was done using CAMPSing (http://www.campsign.bicnirrh.res.in/) (Waghu … Idicula-Thomas, 2020). The probability of the identified peptide sequences to be AMPs was estimated using the Support Vector Machines-VSM algorithm of the CAMP database (Collection of Anti-Microbial Peptides) (http://www.camp3.bicnirrh.res.in/) and DBAASP (Database of Antimicrobial Activity and Structure of Peptides) (https://dbaasp.org/home). A rational design to determine amino acid substitutions that enhance the antimicrobial activity of peptides was done using CAMP’s database rational design tool (Gawde et al., 2022).

### Theoretical Physicochemical Properties

To complement the theoretical information of designed peptides, calculations were done to predict physicochemical properties like a molecular weight in Daltons (Da), net charge, isoelectric point (pI), hydrophobic property, and instability. To achieve this, Protparam Tools through the Expasy server of the UniProt database (https://web.expasy.org/protparam/) (Gasteiger et al., 2005) and Peptide Calculator through Innovagen page were used. The secondary structure was predicted by using PEP-FOLD 3.5 server (http://bioserv.rpbs.univ-paris-diderot.fr/services/PEP-FOLD/) and Pymol viewer.

### Peptide synthesis and characterization

Solid-phase chemical synthesis was performed employing the Fmoc strategy in polypropylene bags according to the methodology described by (Cruz et al., 2014; Guzmán et al., 2020; Prada et al., 2016). The peptides were lyophilized and characterized by high-performance liquid chromatography (HPLC) in a JASCO system (JASCO Corp., Tokyo, Japan), and molecular mass of was determined by electrospray-mass spectrometry (ESI–MS) in a LCMS-2020 ESI–MS equipment (Shimadzu Corp., Kyoto, Japan).

### Reagents and Solvents

All Fmoc-protected amino acids, N,N-diisopropylcarbodiimide (DIC), OxymaPure®, and Fmoc-RinkAmide AM resin (0.6 mmol/g) were purchased from Iris Biotech GmbH (Marktredwitz, Germany). Solvents for synthesis, deprotection reagents, and cleavage reagents used were of synthesis grade and purchased from Merck KGaA (Darmstadt, Germany). Solvents and other chemicals used for high-performance liquid chromatography (HPLC), electrospray ionization (ESI), and matrix-assisted laser desorption/ionization-time of flight (MALDI-TOF) mass spectrometry (MS) and circular dichroism (CD) spectroscopy analyses were of HPLC reagent grade and purchased from Merck KGaA (Darmstadt, Germany) (Guzmán et al., 2020).

### Circular Dichroism Spectroscopy

CD spectroscopy was carried out on a JASCO J-815 CD Spectrometer (JASCO Corp., Tokyo, Japan) in the far ultra-violet (UV) range (190–250 nm), using quartz cuvettes (0.1 cm path length). Each spectrum was recorded averaging three scans in continuous scanning mode. The solvent blank was subtracted from each sample spectrum. Molar ellipticity was calculated for each peptide using 250 µL of 2 mM peptide in 30% (v/v) 2,2,2-trifluoroethanol. The resulting data were analysed using Spectra Manager software (Version 2.0, JASCO Corp., Tokyo, Japan) (Luna et al., 2016).

### Bacterial Strains and Growth Conditions

*Staphylococcus aureus* ATCC 29213, *S. aureus* ATCC 43300, *E. coli* O157:H7, *E. coli* ATCC 25922, and *E. coli* ATCC 43888 strains were purchased from the American Type Culture Collection (ATCC; Rockville, MD, USA). *S aureus* CMPUJ 015 strain was purchased from the Collection of Microorganisms-Pontificia Universidad Javeriana, which is a certified institution belonging to the World Federation of Culture Collection; antibiotic resistance pattern data were presented in the supplementary information (Table S1). Before being used for the antimicrobial assays, Gram positive strains were grown in Müller Hilton broth (MH) at 37 ◦C, and Gram-negative strains were grown in Luria Bertani broth (LB) at 37 ◦C.

### Determination of Minimum Inhibitory Concentration (MIC_50_)

The antimicrobial effect was evaluated by using the broth microdilution method described by CLSI-M07-A10-2015 (CLSI, 2015) adapted for new antimicrobial compounds (Cruz et al., 2017). The evaluation of the minimum inhibitory concentration at 50% of the microbial population (MIC_50_) was determined as follows: a culture of each microorganism was grown in MH for 12 h at 37°C with constant agitation at 200 rpm; these were adjusted until a concentration of 1.5×108 CFU/mL. Then, 100 µL of the inoculum was mixed with 100 µL of mucus fraction in microplates for final concentrations of de 5 µM, 10 µM, 20 µM, 30 µM, 40 µM, and 50 µM. Microplates were incubated at 37°C with constant agitation at 200 rpm. Microbial growth was measured using a Multiskan sky spectrophotometer (Thermo Labsystems Inc., Beverly, MA, USA) at 595 nm every hour for 8 h.

### Determination of cytotoxic and haemolytic activity

Cell viability was determined using the MTT (3-(4,5-dimethylthiazol-2-yl)-2,5-diphenyltetrazolium bromide salt) colorimetric method described by (Mosmann, 1983). Vero (African green monkey kidney epithelial cells) and HaCat (immortalized human epidermal keratinocytes) cells were obtained from ATCC (Saint Cloud, MN, USA). Cells were plated in 96-well plates (5,000 cells/well equivalent to 80% cell confluence) with 200 µL of culture medium. complete (DMEM with 10% SBF and PenStrep 1X) and incubated at 37°C with 5% CO2 and 95% humidity, during 24 hours for complete adherence. Subsequently, lyophilized peptides were dissolved in DMEM culture medium and 200 µL of peptide solutions of 50 µM, 25 µM, 12.5 µM, and 6.25 µM were added using the stock solutions of each peptide as solute and 2% SBF DMEM and 1X PenStrep as solvent. These solutions were incubated with the cells for 48 hours. Additionally, Vero and HaCat cells adhered with 200 µL DMEM at 2% SBF and PenStrep 1X were used as growth control. After MTT addition (500 µg/mL in 1X PBS buffer) lectures were done at 570 nm in a microplate reader (CLARIOstar Plus, BMG Labtech, Ortenberg, Germany).

Haemolysis assay was conducted according to the method described by (Prada et al., 2016). Human erythrocytes were washed 4 times with PBS buffer and suspended at the ratio of 8×109 blood cells/mL. Peptides solutions were added to each test tube at a 1:40 (blood cells: peptide) ratio in a peptide concentration from 120 µM, 60 µM, 30 µM, 15 µM, and 7.5 µM. Samples were incubated for 10 min with constant stirring at room temperature. After incubation, the tubes were centrifuged to separate intact cells and debris. The absorbance of the supernatant obtained at 530 nm against a blank solution (which contained only the diluted sample in PBS) was measured using Multiskan sky spectrophotometer (Thermo Labsystems Inc., Beverly, MA, USA). Each test was conducted in triplicate.

### Data analysis

Unless otherwise stated, graphics and statistical calculations were performed by the GraphPad Prism v6.1 for Windows. (GraphPad software. San Diego. CA. USA). Yield. All the experiments were performed in triplicates and one-way analysis of variance (ANOVA) was used to analyze the differences among the treatments. In all cases, the level of significance was 0.05. The assumption of normality and equality of variances of data was previously tested using Shapiro-Wilk and Levene’s test, respectively.

## Results

### Peptide design

Peptides and their analogs were designed *in silico* seeking to obtain the highest probability of being an AMP. The sequences of the initially designed peptides and their analogs are presented in Table 1. The modifications in the sequences sought to increase the net positive charge, the hydrophobic nature, and the percentage probability of being an antimicrobial peptide, physicochemical properties directly related to the antimicrobial activity (Ortiz López, 2019; Prada et al., 2016).

**Table 1.**
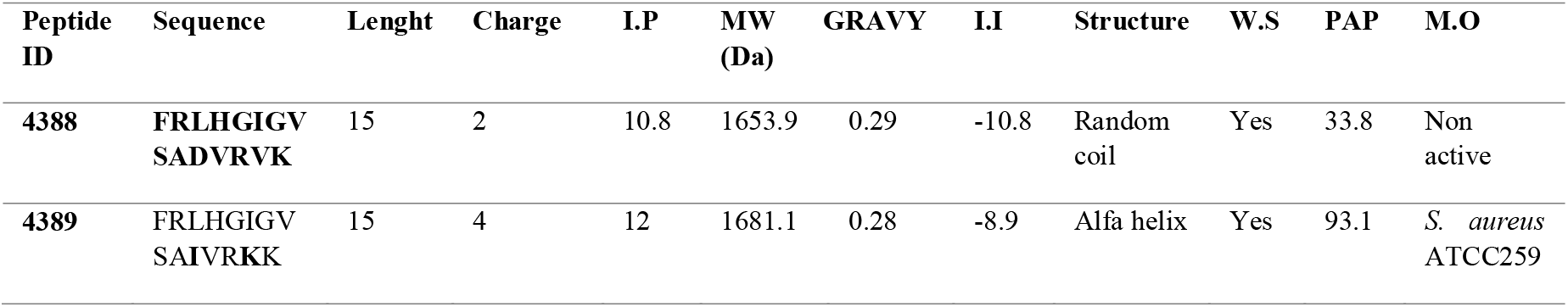
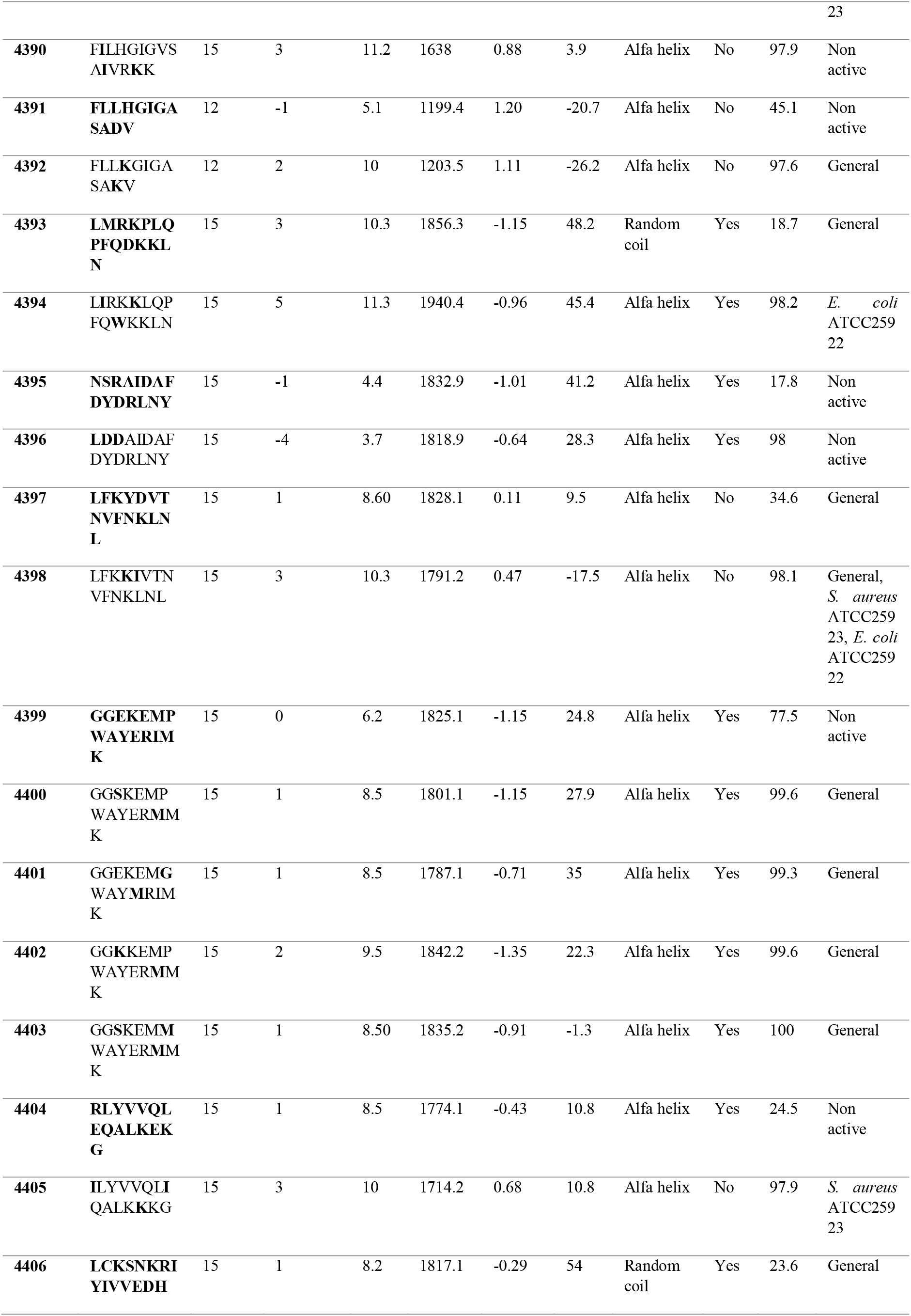
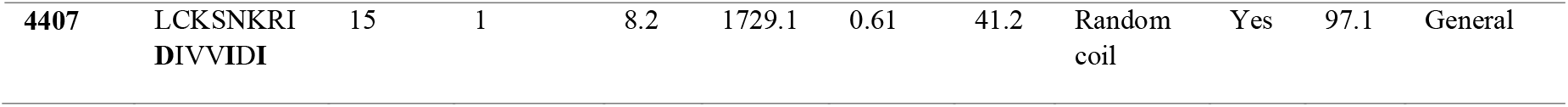
The sequences of the native peptides (identified in the mucous secretion) are indicated in bold. pI: Isoelectric point. GRAVY: hydrophobicity index. Hydrophobic positive, hydrophilic negative. PAP: percentage probability of being an AMP according to the Support Vector Machines algorithm of the CAMP database. M.O: prediction of activity against specific microorganisms according to the DBAASP database.

Peptides were designed *in silico* and visualized with the PEP-FOLD 3.5 program. Figure 1 shows peptides 4397, 4398, 4405, and 4406 that showed the best antimicrobial activity against *S. aureus* strains.

**Figure 1.**
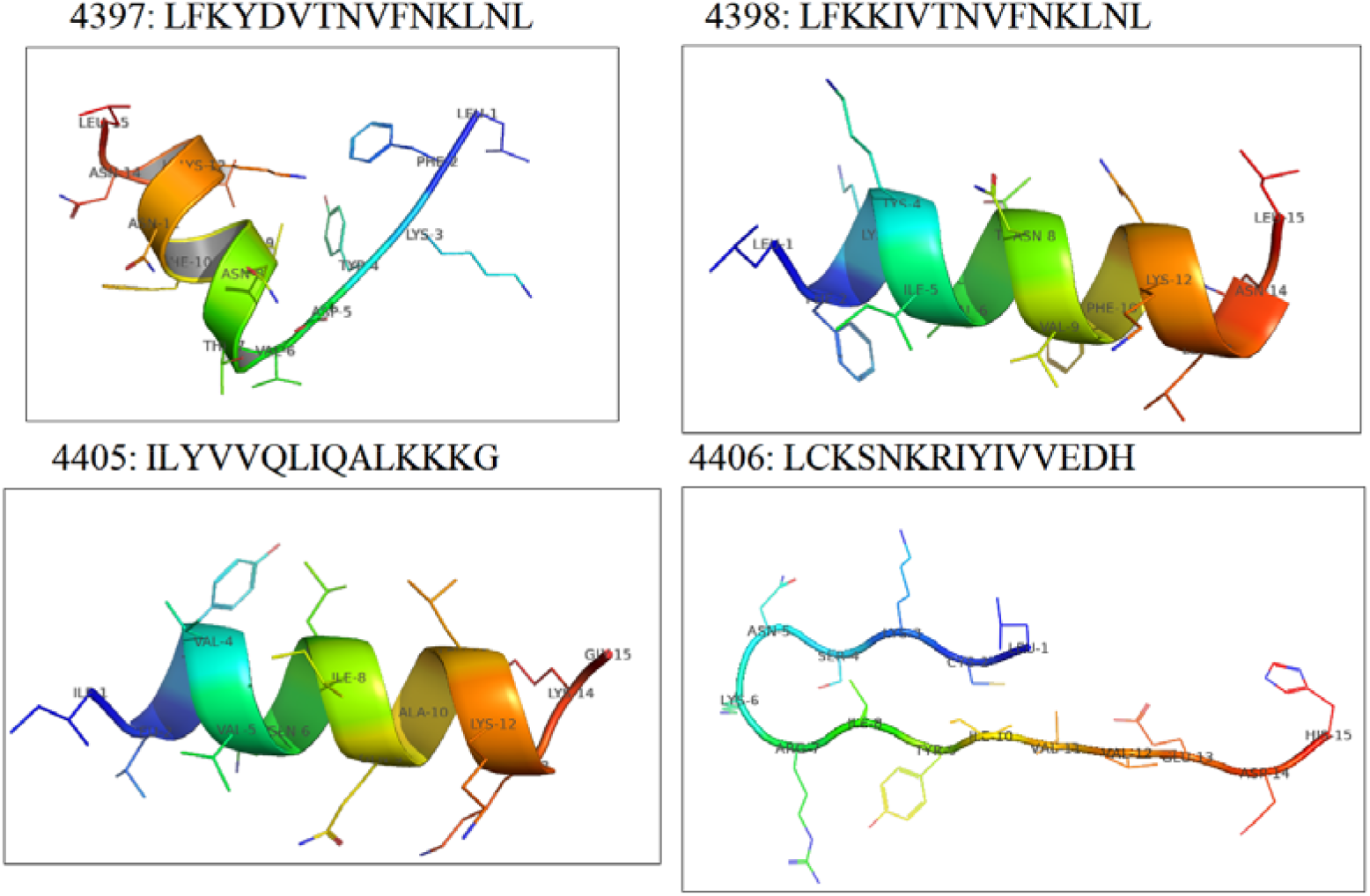
Simulation of the secondary structure of the designed peptides 4397, 4398, 4405, 4406. The construction of the *in silico* (theoretical) secondary structure was carried out in the PEP-FOLD 3.5 database, taking into account the crystallographic structure of the proteins contained in the database, employing sequence homology, using the DEPRAMS algorithm.

### Peptide synthesis and characterization

The 20 peptides reached a degree of purity of approximately 95 to 98% (Table S1). In all cases, it was possible to confirm the sequences by mass analysis. The theoretical and experimental molecular masses of each peptide are presented in Table S1. The results obtained through the analysis with this technique suggest that the peptides were synthesized satisfactorily since the experimental data of mass units (m/z) for each one agreed with their respective theoretically obtained molecular masses.

Regarding the secondary structure of the peptides, the spectrum of circular dichroism was determined for peptides 4397, 4396, 4405, and 4406 under simulated membrane conditions in the presence of TFE/H_2_O at a concentration of 30 % (v/v). The peptides 4397, 4396 and 4405 adopted an alpha-helix structure and exhibited a minimum of two negative bands at 205 and 218 nm, indicating that they adopt a well-defined alpha-helix structure (Figure 2). On the other hand, peptide 4406 showed a strong negative band around 216 nm, a positive band between 195 and 200 nm, and another negative band around 190 nm. These spectra present greater variability than those of an alpha helix, both in amplitude and in the position of the bands. This type of spectrum is associated with beta-sheet structures. This observation disagrees with the three-dimensional model that predicts a peptide with a random coil structure. The typical spectrum of a random coil structure is characterized by an intense negative band around 200 nm and another very weak band, which can be positive or negative, around 220-230 nm (Carvajal-Rondanelli et al., 2018) (Figure 2).

**Figure 2.**
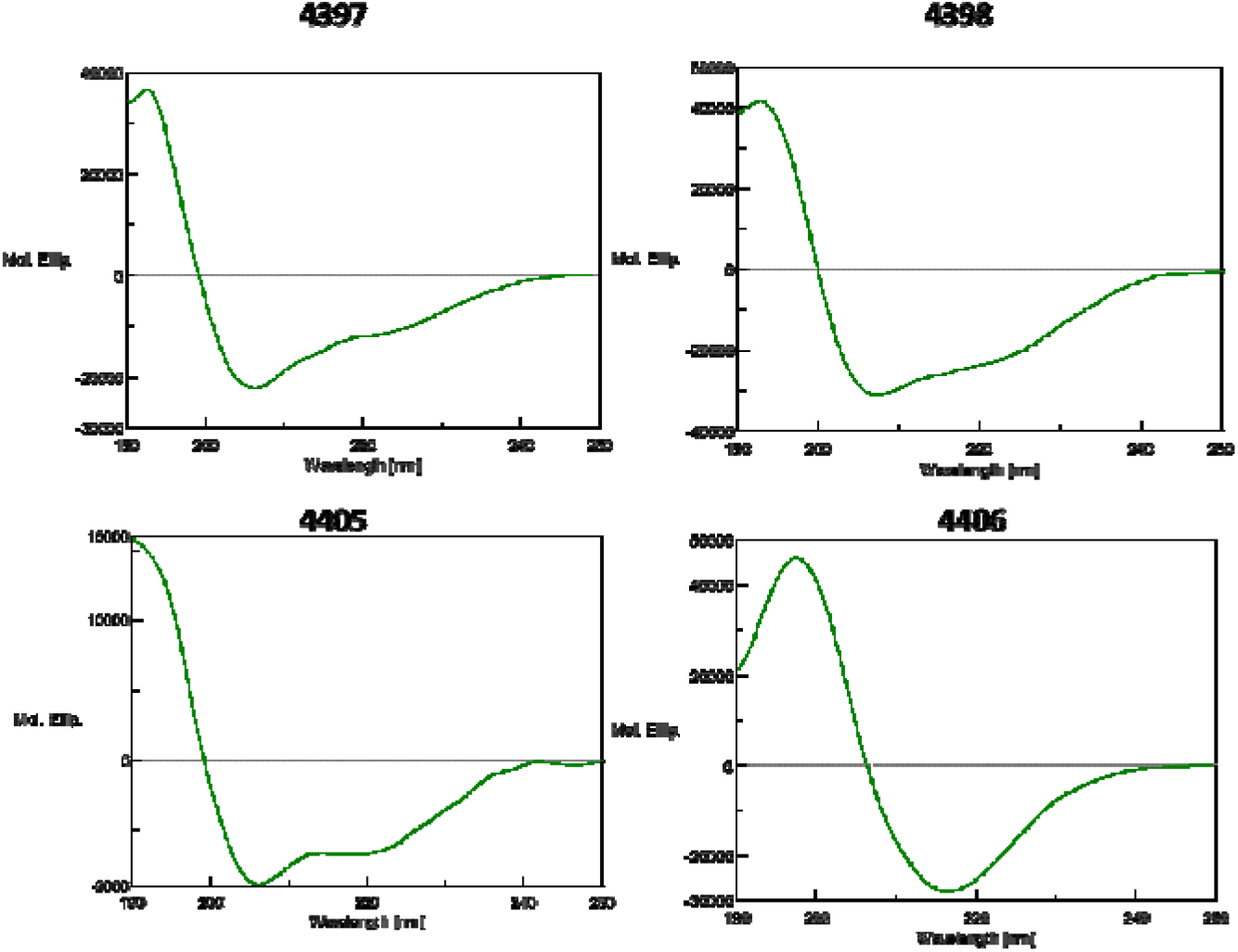
Circular dichroism spectra of peptides 4397, 4398, 4405, and 4406.

### Antimicrobial activity assay

18 of the 20 synthetized peptides inhibited the growth of *S. aureus* strains at a concentration below 50 μM. For *E. coli* strains no inhibitory activity was observed. Ofloxacin was used as a positive control (Table 2). The peptides with the greatest antibacterial activity against *S. aureus* strains were 4397, 4398, 4405, and 4406 with an MIC_50_ ranging from 20 to 50 μM (Table 3).

**Table 2.**
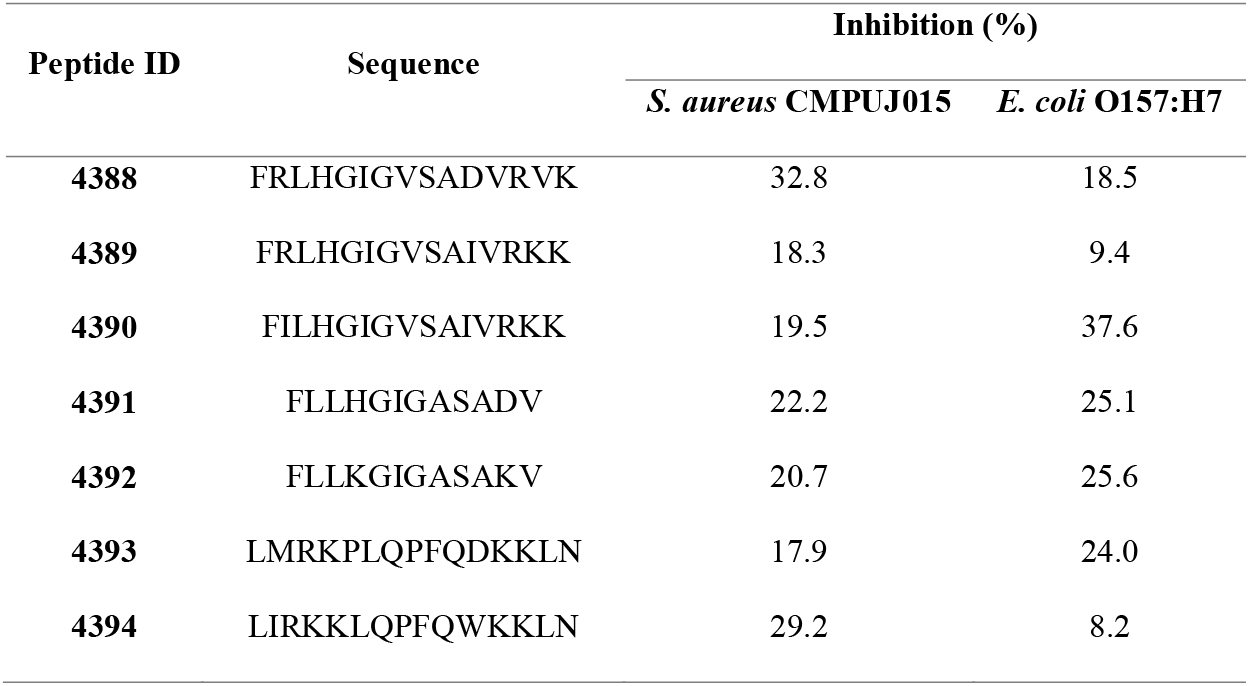
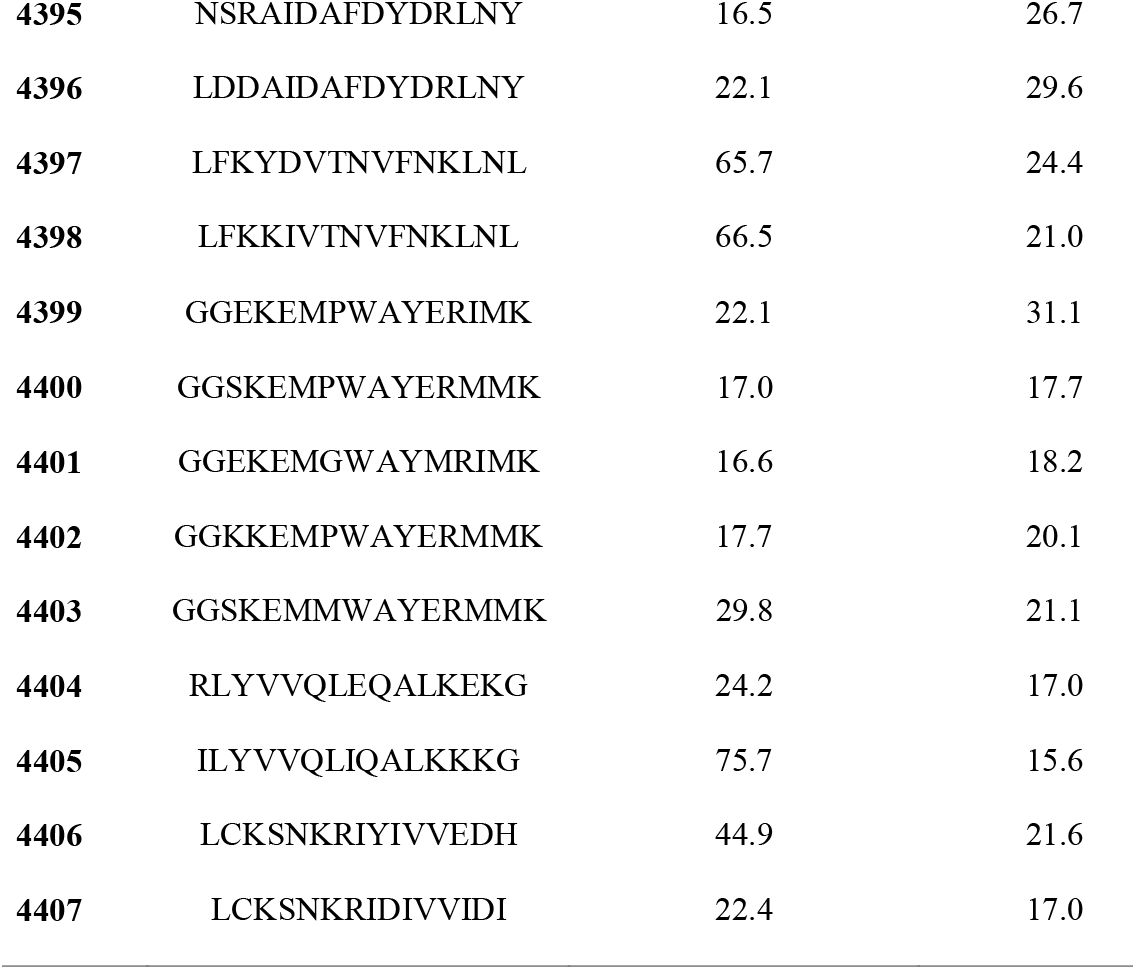
Screening antimicrobial activity of the peptides at 50 µM against *S. aureus* CMPUJ015 and *E. coli* O157:H7. The assays were performed in triplicate and the results are the means of three values ± standard deviation. For these assays, a growth control was used.

**Table 3.**
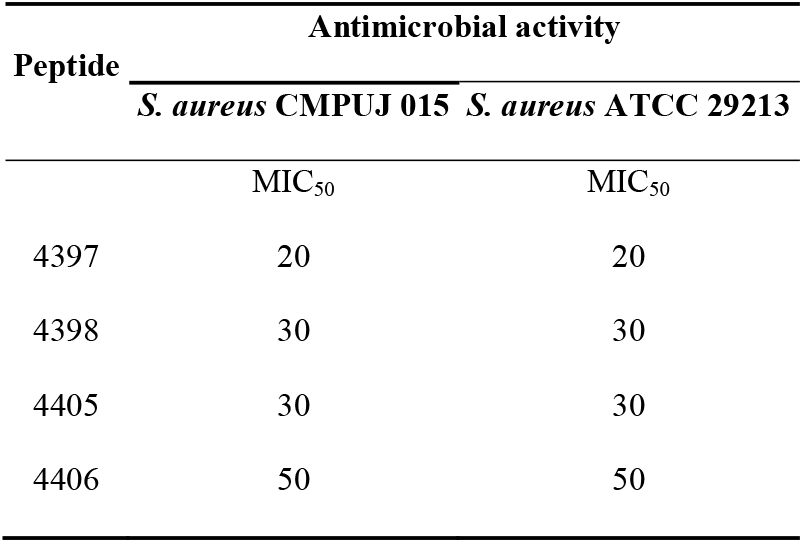
Minimum Inhibitory Concentration 50 (MIC_50,_ µM) of the peptides 4397, 4398, 4405 and 4406 on *S. aureus* CMPUJ 015 and *S. aureus* ATCC 29213.

### Cytotoxic and haemolytic activity assay

Peptides 4397, 4398, and 4405 showed no toxic effect on Vero and HaCat cells at concentrations two times CMI_50_. CC50 was not obtained since for all evaluated peptides it turns out to be significantly greater than 50 µM. Therefore, an exact value of IS could not be calculated. To determine the IS, the concentration of the peptides would have to be increased to values well above the MIC_50_. On the other hand, little or no haemolysis was detected at concentrations of 120 µM, while peptide 4405 had haemolytic activity of 34.5% at 60 µM (Table 4).

**Table 4.**
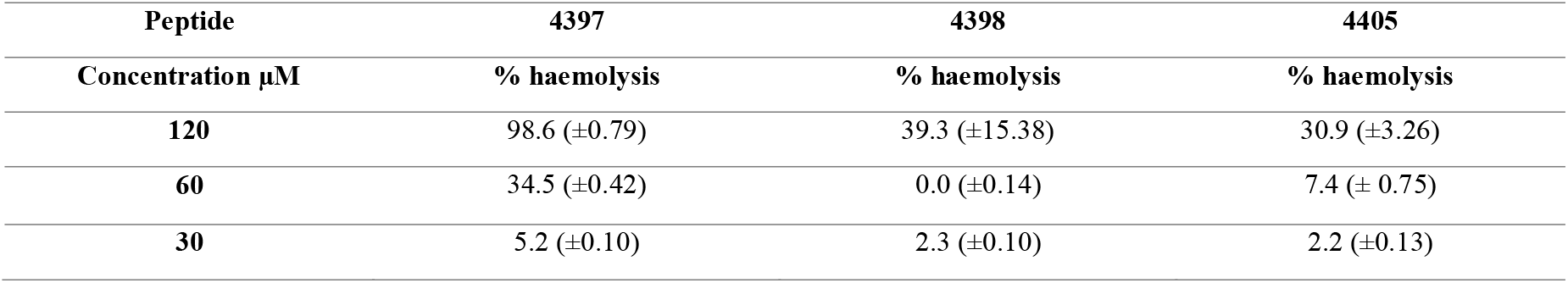
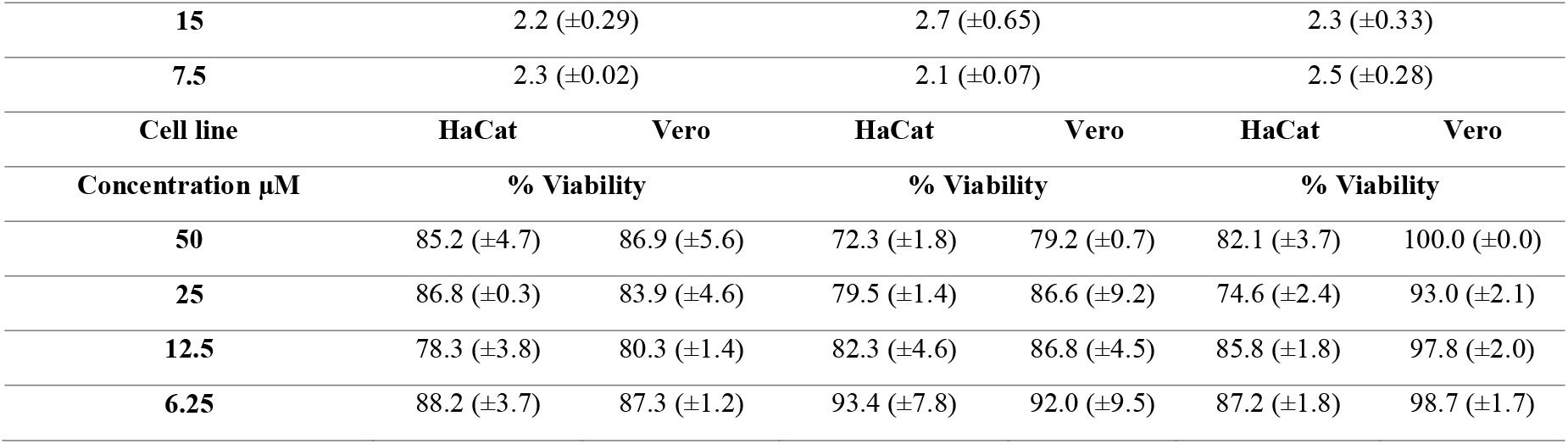
Haemolytic and cytotoxic activity of peptides. Assays were performed in triplicate and the results are the mean of three values ± standard deviation.

| Peptide | 4397 | 4398 | 4405 |
| --- | --- | --- | --- |
| Concentration µM | % haemolysis | % haemolysis | % haemolysis |
| **120** | 98.6 (±0.79) | 39.3 (±15.38) | 30.9 (±3.26) |
| **60** | 34.5 (±0.42) | 0.0 (±0.14) | 7.4 (± 0.75) |
| **30** | 5.2 (±0.10) | 2.3 (±0.10) | 2.2 (±0.13) |

|  |  |  |  |  |  |  |
| --- | --- | --- | --- | --- | --- | --- |
| 15 | 2.2 (±0.29) |  | 2.7 (±0.65) |  | 2.3 (±0.33) |  |
| 7.5 | 2.3 (±0.02) |  | 2.1 (±0.07) |  | 2.5 (±0.28) |  |
| Cell line | HaCat | Vero | HaCat | Vero | HaCat | Vero |
| Concentration μM | % Viability |  | % Viability |  | % Viability |  |
| 50 | 85.2 (±4.7) | 86.9 (±5.6) | 72.3 (±1.8) | 79.2 (±0.7) | 82.1 (±3.7) | 100.0 (±0.0) |
| 25 | 86.8 (±0.3) | 83.9 (±4.6) | 79.5 (±1.4) | 86.6 (±9.2) | 74.6 (±2.4) | 93.0 (±2.1) |
| 12.5 | 78.3 (±3.8) | 80.3 (±1.4) | 82.3 (±4.6) | 86.8 (±4.5) | 85.8 (±1.8) | 97.8 (±2.0) |
| 6.25 | 88.2 (±3.7) | 87.3 (±1.2) | 93.4 (±7.8) | 92.0 (±9.5) | 87.2 (±1.8) | 98.7 (±1.7) |

## Discussion

In recent years, haemocyanins, copper-containing respiratory proteins found in many mollusks, have emerged as a potential source of AMPs. *A. fulica* haemocyanin (AfH) has not been thoroughly explored for its antimicrobial potential. To our knowledge, studies of hemocyanin from marine gastropods as a source of antibacterial AMPs have been studied since 2015, when Zhuang et al. described haliotisin, a region of abalone hemocyanin that appears to perform an anti-infective function. This region is a highly conserved region amongst molluscan hemocyanin and synthetic peptides designed on the amino acid composition of this region target both Gram-positive and Gram-negative bacteria (Zhuang et al., 2015).

Regarding terrestrial gastropods, in 2020, Dolashki et al. discovered three potential AMPs in the mucous secretion of the *Cornu aspersum*. These AMPs exhibited high homology with hemocyanins isolated from snails *Helix aspersa, Helix pomatia*, and *Helix lucorum* (Dolashki et al., 2020). It is hypothesized that proteolytic processes may have played a role in the generation of these peptides within the mucous secretion. Notably, some of the identified peptides displayed elevated levels of glycine, leucine, and proline residues, suggesting their potential importance in maintaining the stability of their antimicrobial activity.

In this study, 15-residue synthetic peptides with alpha-helical structures and cationic properties exhibited the highest biological activity against Gram-positive strains, with MIC values ranging from 20 to 30 µM. Contrary to bioinformatics predictions, peptides 4397 (AfH5) and 4398 (AfH5-4K5I) showed similar inhibition percentages against both the Gram-positive strain *S. aureus* CMPUJ015 (65.73% and 66.47% inhibition percentages, respectively), and the Gram-negative *E. coli* O157:H7 (inhibition percentages 6.1% and 5.1%, respectively). Theoretical prediction of antimicrobial activity suggested that peptide 4398 had a higher probability of having an antimicrobial activity (98.1%) than peptide 4397 (34.6%).

The support Vector Machines (SVM) algorithm of the CAMP database uses amino acid composition, hydrophobicity, net charge, peptide length, and secondary structure to predict the antimicrobial activity of a peptide (Wang et al., 2009, 2016). The difference in the antimicrobial activity probability between peptide 4397 and its analog 4398 may be due to the net positive charge of +3 of peptide 4398 compared to +1 for peptide 4397. The net charge of an antimicrobial peptide is an important property that influences its antimicrobial activity. In general, antimicrobial peptides with a net positive charge have been found to have higher antimicrobial activity than those with a net neutral or negative charge (Hale … Hancock, 2007; Huang et al., 2010; Stark et al., 2002). Furthermore, peptide 4398 incorporates a lysine residue at position 4 instead of the tyrosine present in peptide 4397. Lysine is a positively charged amino acid that can interact with the cell membranes of microorganisms and enhance the antimicrobial activity of the peptide (Nguyen et al., 2011). Peptide 4398 also includes an isoleucine residue at position 5 in place of aspartic acid. Isoleucine is a hydrophobic amino acid that may be involved in antimicrobial activity by increasing the affinity of the antimicrobial peptide for bacterial membranes (Fjell et al., 2012). Therefore, in this case, the predictions made by bioinformatics tools did not match the experimental results.

Although the criteria used by the CAMP database SVM algorithm to predict the antimicrobial activity of a peptide may be useful to identify new peptides with potential antimicrobial activity, the experimental evaluation of antimicrobial activity is essential. to validate these predictions. The antimicrobial activity of a peptide does not only depend on the presence of certain amino acids but also on the specific sequence and three-dimensional structure of the peptide.

Even though there are studies that demonstrate the antimicrobial properties of the mucus secretion of *A. fulica*, little is known about the activity oofresistant bacteria. Our previous studies confirmed the inhibitory action of mucoprotein and protein fractions of mucus secretion against *S. aureus* CMPUJ 015 (Pereira et al., 2016; Suárez et al., 2021). In this study, new analogs of haemocyanin-derived peptides identified in >30kDa protein fractions from mucus secretion of *A. fulica*, were also active against *S. aureus* CMPUJ 015, *S. aureus* ATCC 29213, *S. aureus* ATCC 43300 and *S. aureus* ATCC 25922.

Evaluated peptides were found to exhibit a bacteriostatic effect rather than a bactericidal effect on *S. aureus* strains. This conclusion is drawn based on the observation of increased culture absorbance after 24 hours of exposure to the fractions, indicating the growth of viable cells (data not shown). However, additional tests are required to further investigate the mechanism of action of AfH-derived peptides.

No inhibitory activity of AfH-derived peptides was found against *E. coli* O157:H7. AMPs active against Gram-positive bacteria and those active against Gram-negative bacteria differ in several aspects, including amino acid composition, structure, and electrical charge (Brtz-Oesterhelt … Sass, 2010). In general, peptides active against Gram-positive bacteria tend to have a net positive charge and a more rigid structure, whereas peptides active against Gram-negative bacteria tend to have a net negative charge and a more flexible structure (Brtz-Oesterhelt & Sass, 2010; Hancock & Sahl, 2006; Li et al., 2012; Pasupuleti et al., 2012). This is due to the characteristics of the outer membrane of Gram-negative bacteria, which is composed of a lipid bilayer and a lipopolysaccharide layer that acts as an additional barrier for the penetration of antimicrobial compounds (Nikaido, 2003). Based on the above, it is possible that, in this case, the net charge is not the main reason that contributes to explaining the lower inhibitory effect against *E. coli* O157:H7 compared to *S. aureus* CMPUJ015.

Selected peptides (4397, 4398, and 4405) with the best antimicrobial activity did not present cytotoxic or haemolytic effects against the evaluated cell lines. However, further research is needed to determine the cytotoxic effects of these peptides on different types of human cells and to better understand the mechanisms underlying their cytotoxic activity.

## Conclusions

Our results highlight the potential of *A. fulica* hemocyanin as a valuable protein model for the rational design of novel AMPs targeting bacteria of public health significance. While our study focuses on the antimicrobial activity against Gram-positive strains, further investigations are warranted to evaluate the efficacy of these PAMs against Gram-negative bacteria. The development of effective AMPs derived from *A. fulica* hemocyanin holds promise as a potential therapeutic strategy against antibiotic-resistant bacterial infections, contributing to the ongoing battle against the rising global health threat posed by multidrug-resistant pathogens.

