## Supplementary material for "*Achatina fulica* Haemocyanin-Derived Peptides as Novel Antimicrobial Agents": Table S1

**Table S1.** Antibiogram of *S. aureus* strains. The clinical tests were carried out by the service of Merced Lab, Bucaramanga, Colombia.

| **Antibiotic** | ***S. aureus* CMPUJ 015** | | ***S. aureus* ATTC 29213** | |
|---|---|---|---|---|
| | **MIC (ug/mL)** | **Interpretation** | **MIC (ug/mL)** | **Interpretation** |
| Amox/A Clav | <=4/2 | S | <=4/2 | S |
| Amp/Sulbactam | >16/8 | R | 16/8 | I |
| Ampicillin | >8 | R | >8 | R |
| Cefazolin | <=4 | S | <=4 | S |
| Ciprofloxacin | <=1 | S | <=1 | S |
| Clindamycin | 1 | I | 1 | I |
| Daptomycin | 1 | S | 1 | S |
| Erythromycin | >4 | R | 2 | I |
| Gentamicin | <=4 | S | <=4 | S |
| Induction Clindamycin | <=4/0.5 | NEG | <=4/0.5 | NEG |
| Levofloxacin | <=1 | S | <=1 | S |
| Linezolid | 4 | S | 4 | S |
| Moxifloxacin | <=0.5 | S | <=0.5 | S |
| Nitrofurantoin | <=32 | | <=32 | |
| Oxacillin | 2 | S | 1 | S |
| Penicillin | >8 | R | >8 | R |
| Rifampicin | <=1 | S | <=1 | S |
| Cefoxitin Screening | <=4 | NEG | <=4 | NEG |
| Synercid | <=1 | S | <=1 | S |
| Tetracycline | >8 | R | >8 | R |
| Trimet/Sulfa | <=0.5/9.5 | S | <=0.5/9.5 | S |
| Vancomycin | 2 | S | 2 | S |

S: sensible; I: intermediate sensitivity; R: resistant; NEG: Negative.
